# Phylogenomics and biogeography of the feather lice (Phthiraptera: Ischnocera) of parrots

**DOI:** 10.1101/2023.10.27.564336

**Authors:** Kevin P. Johnson, Jorge Doña

## Abstract

Avian feather lice (Phthiraptera: Ischnocera) have undergone morphological diversification into ecomorphs based on the mechanism for escaping host preening defenses. Parrot lice are one prominent example of this phenomenon, with wing, body, or head louse ecomorphs occurring on various groups of parrots. Currently defined genera of parrot lice typically correspond to this ecomorphological variation. Here we explore the phylogenetic relationships among parrot feather lice by sequencing whole genomes and assembling a target set of 2,395 nuclear protein coding genes. Phylogenetic trees based on concatenated and coalescent analyses of these data reveal highly supported trees with strong agreement between methods of analysis. These trees reveal that parrot feather lice fall into two separate clades that form a grade with respect to the *Brueelia*-complex. All parrot louse genera sampled by more than one species were recovered as monophyletic. The evolutionary relationships among these lice showed evidence of strong biogeographic signal, which may also be related to the relationships among their hosts.

## Introduction

Convergent evolution in morphology can obscure phylogenetic relationships, because distantly related organisms appear similar in form (Futuyma 1998, Bels and Russell 2023). The feather lice (Phthiraptera: Ischnocera) of birds are one prominent example of convergent evolution in insects (Johnson et al. 2012). These wingless ectoparasites spend their entire lifecycle on the body of the host (Clayton et al. 2015), such that hosts are analogous to islands for feather lice. Across the diversity of avian hosts, feather lice have repeatedly diverged in morphology related to how these parasites escape host preening defenses (Clay 1949, 1951). Even though this divergence typically occurs between lice living on the same group of birds (Johnson et al. 2012), across the entire diversity of feather lice, this has led to overall convergence, because there are a limited number of ways in which lice escape host preening defenses.

Wing lice insert between the feather barbs of the wing, where they are difficult for the host to remove by preening with the bill (Clayton et al. 2003). They have a long and slender body form adapted for this insertion. Head lice remain relatively sessile on the feathers of the head, gripping the feather barbs with their mandibles to avoid being removed by scratching, since a bird cannot preen its own head with the bill. This ecomorph has a rounded body and triangular head, with expanded temples housing strong mandibular muscles (Clay 1951). Body lice burrow in the downy portions of the body feathers to escape host preening (Clayton 1991). These lice have a rounded head margin and rounded body form. All three of these ecomorphs occur on parrots and relatives (Aves: Psittaciformes). Previous work has indicated that the various genera of parrot lice are closely related (Johnson et al. 2012, de Moya et al. 2019, de Moya 2022), suggesting that an adaptive divergence of feather lice occurred within this single group of birds. However, given that genera of parrot lice are most notably differentiated based on characters related to ecomorphology (Price et al. 2003), the possibility exists that convergence may further obscure relationships and generic boundaries in this group. Thus, it is important to use additional molecular data sets, with expanded taxon sampling, to uncover the phylogenetic tree of this group of parasites.

The various genera of parrot lice also have intriguing patterns of biogeographic distribution and host association (Price et al. 2003). In the New World, only a single genus (*Paragoniocotes*) is found (Guimarães 1975), and is of the body louse ecomorph. In the Old World, the situation is more complex, with most groups of hosts harboring louse genera of two distinct ecomorphs. The only body louse genus in the Old World (*Psittoecus*) is known only from cockatoos (Guimarães 1974a), which are restricted to Australasia. The wing louse genus *Neopsittaconirmus* occurs broadly on Old World parrots and cockatoos, while the wing louse genus *Psittaconirmus* is restricted to lorikeets, and a few other small-bodied Australasian species, such as fig-parrots (Guimarães 1974b). The three genera of head lice tend to show restricted distributions, both with respect to biogeography and host group. The genus *Echinophilopterus* occurs mainly on African and Asian parrots, but with some occurrence on Australasian hosts, and even a species occurring on a ground-roller (Coraciiformes: *Uratelornis chimaera*) in Madagascar (Guimarães 1980). The head louse genus, *Forficuloecus*, has a more exclusively Australasian distribution, occurring on parrots in Australia and New Zealand (Guimarães 1974a, 1985a, Price et al. 2008). Finally, the head louse genus *Theresiella* is only known from the tiger-parrots (*Psittacela*) of New Guinea (Guimarães 1985b).

Some prior phylogenies of broader feather louse relationships included limited sampling of parrot lice, typically a single species per genus, often with not all genera included (Johnson et al. 2012, de Moya et al. 2019, de Moya 2022). These phylogenies have generally confirmed that parrot lice are closely related to the *Brueelia*-complex, a diverse group of lice that occurs on a variety of avian orders, particularly songbirds (Passeriformes). However, these phylogenies have differed in whether parrot lice are a monophyletic group, or perhaps a paraphyletic clade in which the *Brueelia*-complex is embedded, leaving this as an unresolved question. Given the broad biogeographic and host distribution of parrot lice, it is important to increase taxon sampling for a more complete picture of the evolutionary history of this group. The interplay between ecomorphology, biogeography, and host association remains to be explored.

Here we sample a broad diversity of the feather lice of parrots, including representatives of all genera. We also include broad sampling of the *Brueelia*-complex and outgroup genera. With these samples, we sequence the entire genome using short read shotgun sequencing. From these raw reads, we assemble 2,395 single copy ortholog nuclear genes and use these sequences to reconstruct a phylogenomic tree of parrot lice and relatives.

## Materials and Methods

### Taxon sampling

Samples of 54 individuals of parrot feather lice were included for genome sequencing (Table 1). These were selected from material stored in 95% ethanol at –80C. This taxon sample included representatives of all seven genera of parrot feather lice. Six of these genera (all but *Theresiella*) were sampled by more than one species. In some cases, multiple individuals of the same louse species, but from different host species were included because of the possibility of cryptic species. As additional potential ingroups, we selected 25 samples (Table 1) from among genera of the *Brueelia*-complex that have been shown by prior studies (de Moya et al. 2019), to be closely related to parrot feather lice. In addition, we selected 17 additional more distantly related outgroup taxa (Table 1) from among the *Quadraceps*-complex, *Philopterus*-complex, *Goniodes*-complex, and *Heptapsogaster*-complex, among others, to serve to root the tree based on higher level phylogenies of avian feather lice (de Moya et al. 2019).

**Table 1.**
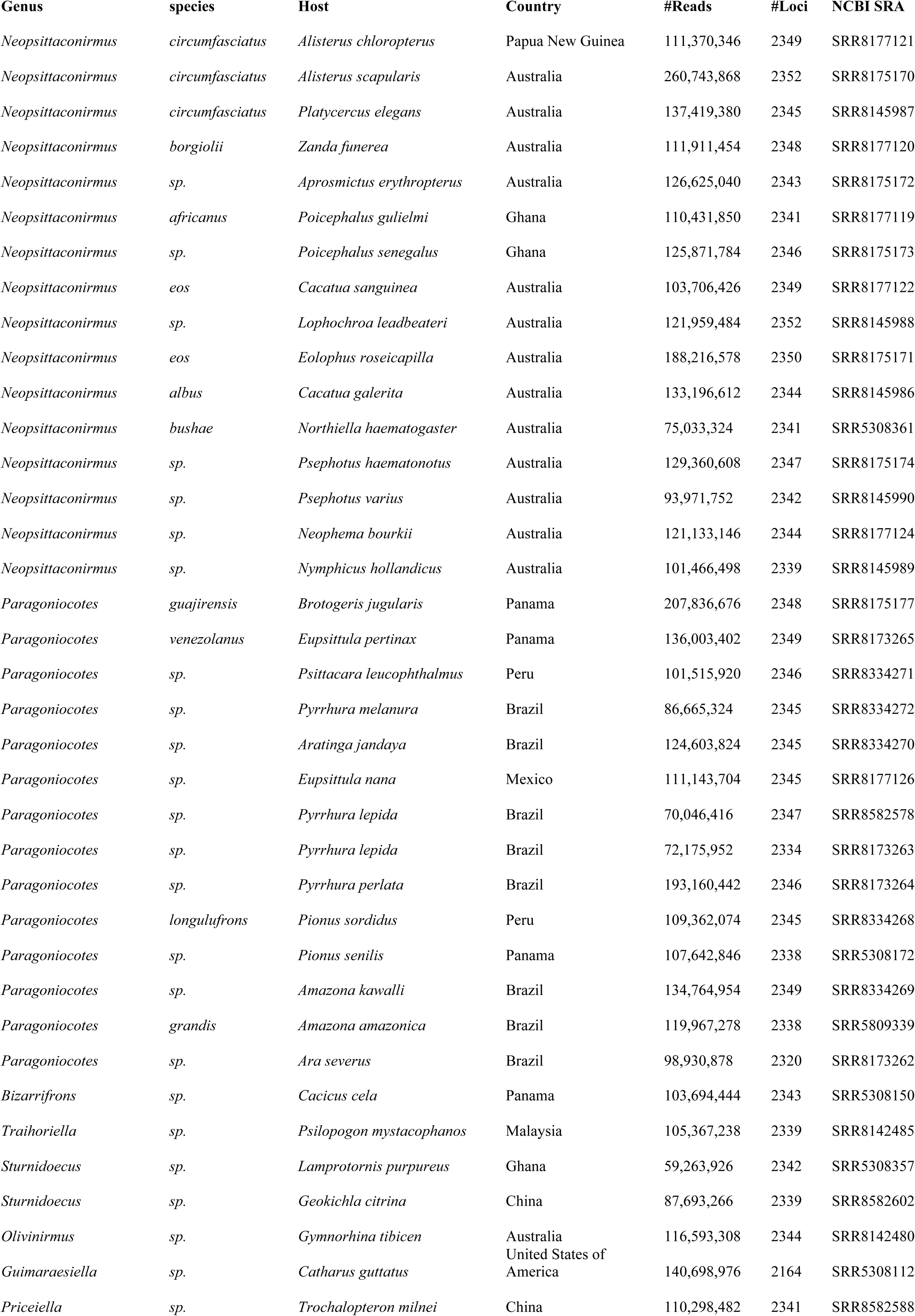
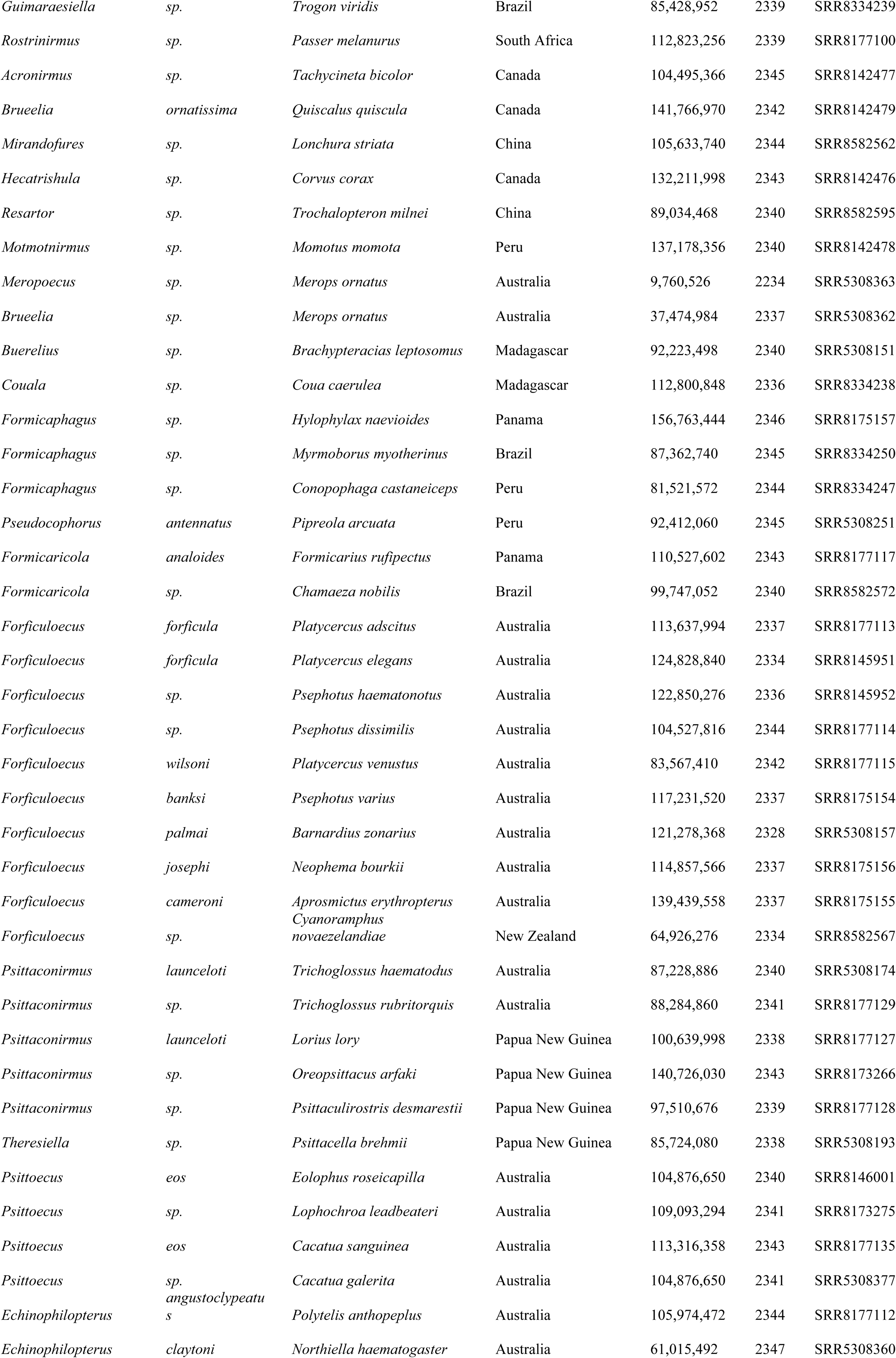
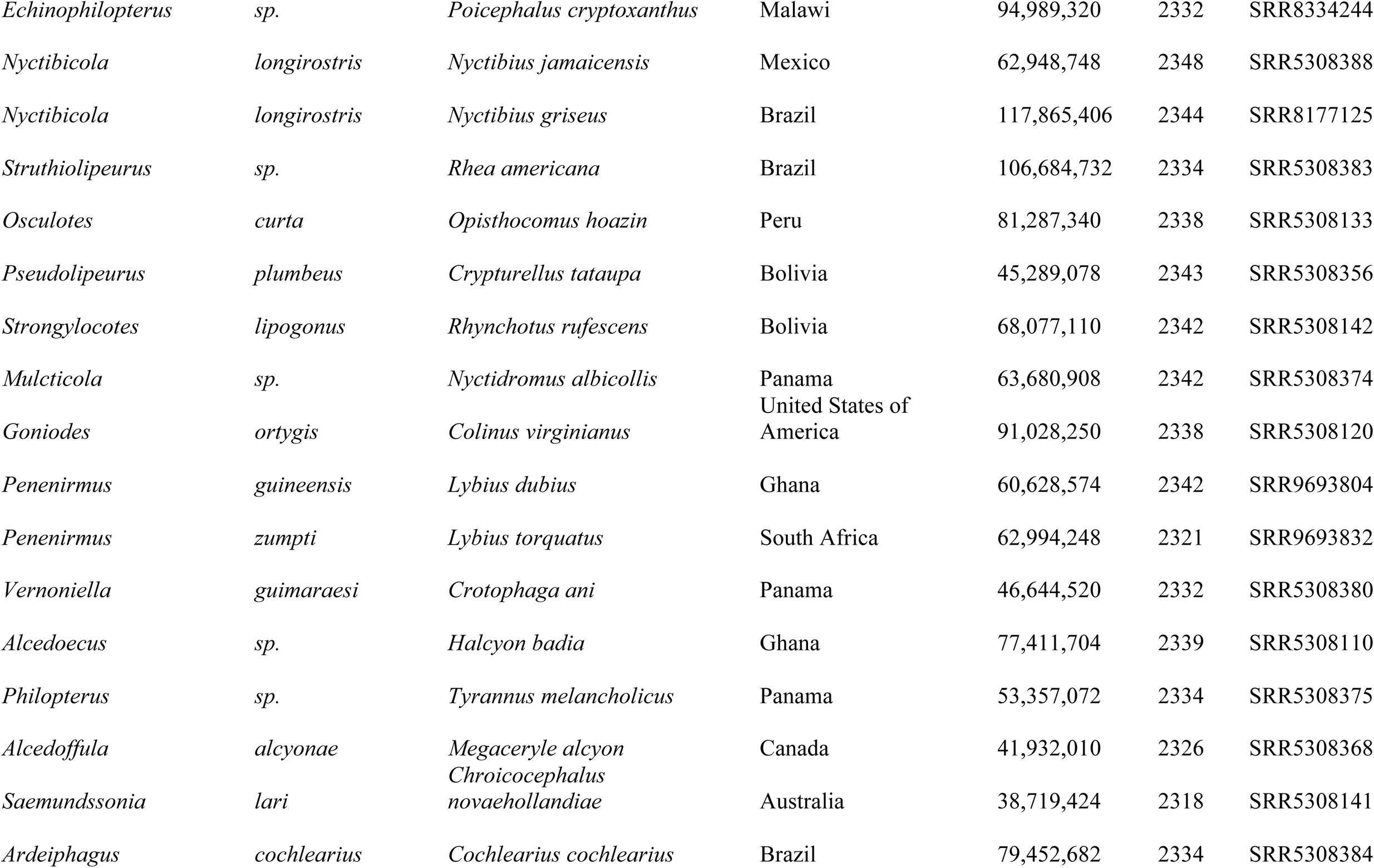
Samples in study.

### DNA extraction and sequencing

Prior to DNA extraction, each specimen was photographed as a voucher (https://figshare.com/s/4f33cbe035e9da9b846c; 10.6084/m9.figshare.23807646). Lice were identified to the genus level using Price et al. (2003), and tentative species assignment was made based on prior determination of slide mounted specimens from the same lot (Price et al. 2008) or from prior known host associations (Price et al. 2003). Total genomic DNA was extracted by grinding the specimen and then using a Qiagen QIAamp DNA Micro Kit (Qiagen, Valencia, CA, USA) for the extraction procedure. The initial incubation with proteinase K was performed for 48 hours, but otherwise manufacturer’s protocols were followed. From this extraction (50ul in buffer AE), a Hyper library construction kit (Kapa biosystems) was used to prepare paired end libraries of around 400-500 bp inserts. These libraries were sequenced using 150bp paired-end reads on an Illumina NovaSeq 6000 machine with S4 reagents. Libraries were multiplexed (typically 48 per lane) to achieve around 30-60X coverage of the nuclear genome, assuming a 200Mbp genome size. The software bcl2fastq was used to trim adapters and demultiplex the reads into fastq files. We deposited the raw reads from each library on the NCBI SRA (Table 1).

### Phylogenomic analyses

Additional adaptor and quality trimming was performed using *fastp* v0.20.1 (Chen et al. 2018). These libraries were then used in the aTRAM 2.0 (Allen et al. 2018) pipeline to assemble 2,395 single-copy ortholog genes (Johnson et al. 2018) using reference amino acid sequences from the human louse (*Pediculus humanus*). The aTRAM assemblies used the ABySS assembler, with *tblastn* searches of three iterations and max-target-seqs set to 3,000. From the assembled contigs for each gene, the Exonerate (Slater and Birney 2005) pipeline of aTRAM (atram_stitcher.py) was used to identify exon sequences and stitch them together in a single contig. The gene sequences were then aligned using MAFFT (Katoh et al. 2002) and trimmed according to procedures detailed in Johnson et al. (2021).

Gene alignments were used in both concatenated and coalescent phylogenomic analyses. For the concatenated alignment, IQ-TREE 2 (Minh et al. 2020) was used to identify the optimal number of partitions. Using these partitions and the model parameters estimated for each of them, we performed optimal tree searches and conducted bootstrapping using the ultrafast option. For the coalescent analysis, IQ-TREE 2 was used to reconstruct individual gene trees for each gene alignment. These gene trees were combined into a coalescent species tree with local posterior probabilities for each node using ASTRAL III (Zhang et al. 2018).

To explore the possibility of cryptic species, such that only a single individual of each species was included in the biogeographic reconstruction, we used the genomic sequencing libraries to assemble the mitochondrial COI gene. For each library, we subsampled four million total reads (two million read 1 and two million read 2) using Seqtk v 1.3. As a starting point, we used a partial COI sequence from *Forficuloecus cameroni* (Price, Johnson & Palma, 2008). We then used aTRAM 2.0 (Allen et al. 2018) to assemble reads from the same species with the goal of extending this partial sequence to obtain a complete COI sequence. After validating the full sequence and confirming the presence of a complete open reading frame, this sequence was then employed as the reference for the assembly of all other samples in our study.

The COI sequences were aligned based on amino acids using MAFFT. From this alignment, we estimated a phylogenetic tree using ML searches in IQ-TREE 2, with ulstrafast bootstrapping. Raw uncorrected sequence divergences were calculated for all pairwise comparisons using R (*dist.dna* from APE v5.5, Paradis and Schliep 2018). We plotted these comparisons for each species, and examined these plots for substantial gaps. Because a 5% sequence divergence cutoff has been used for prior studies of lice (Johnson et al. 2021), we elected to use this cutoff for the purposes of delimiting terminal taxa for biogeographic reconstruction.

For biogeographic reconstruction, we used BioGeoBears v1.1.2 (Matzke 2013, 2014). Because this approach requires and ultrametric tree, we performed a dating analysis in IQ-TREE 2 using the least square dating method (To et al. 2016). We set the root age at 34 mya based on de Moya (2021). For additional calibration, we identified five terminal cospeciation events using eMPRess v1.2.1 (Santichaivekin et al. 2020). These events involved terminal sister species of lice associated with terminal sister species of hosts. For these calibration points, we used confidence intervals obtained from TimeTree or single estimates where only as a single study was available (Kumar et al. 2022). Further details on these calibration points are provided in Table S1. We set a minimum branch length constraint of 0.01 and calculated confidence intervals. Our approach aligns with methodologies on dating analyses implemented in prior louse phylogenomic studies (Johnson et al. 2021, 2022). Using this ultrametric louse tree, we ran BioGeoBears for the models DEC, DIVALIKE, and BAYAREALIKE. Each model was also run an additional time with the parameter J, to allow for jump dispersal events (Matzke 2014). The model with the lowest AIC score was then used to estimate ancestral ranges over the louse tree.

## Results

### Higher level phylogeny

Of the 2,395 single copy ortholog genes targeted for assembly, between 2,164 and 2,352 of these were assembled with aTRAM depending on the sample (Table 1). A total of 2,370 genes passed the filter for inclusion, and after trimming the alignment the concatenated matrix was 3,914,892 aligned bp. The partitioned concatenated IQ-TREE analysis produced a highly resolved tree, with 76 out of 78 ingroup nodes supported by 100% ultrafast bootstrap replicates (Figure 1). The ASTRAL coalescent analysis also produced a highly resolved tree, with 77 out of 78 ingroup nodes with 1.0 local posterior probability (Figure 1 and Supplemental Figure 1). The concatenated and coalescent trees were identical except for two local branch rearrangements, one in the ingroup and one in the outgroup.

**Figure 1.**
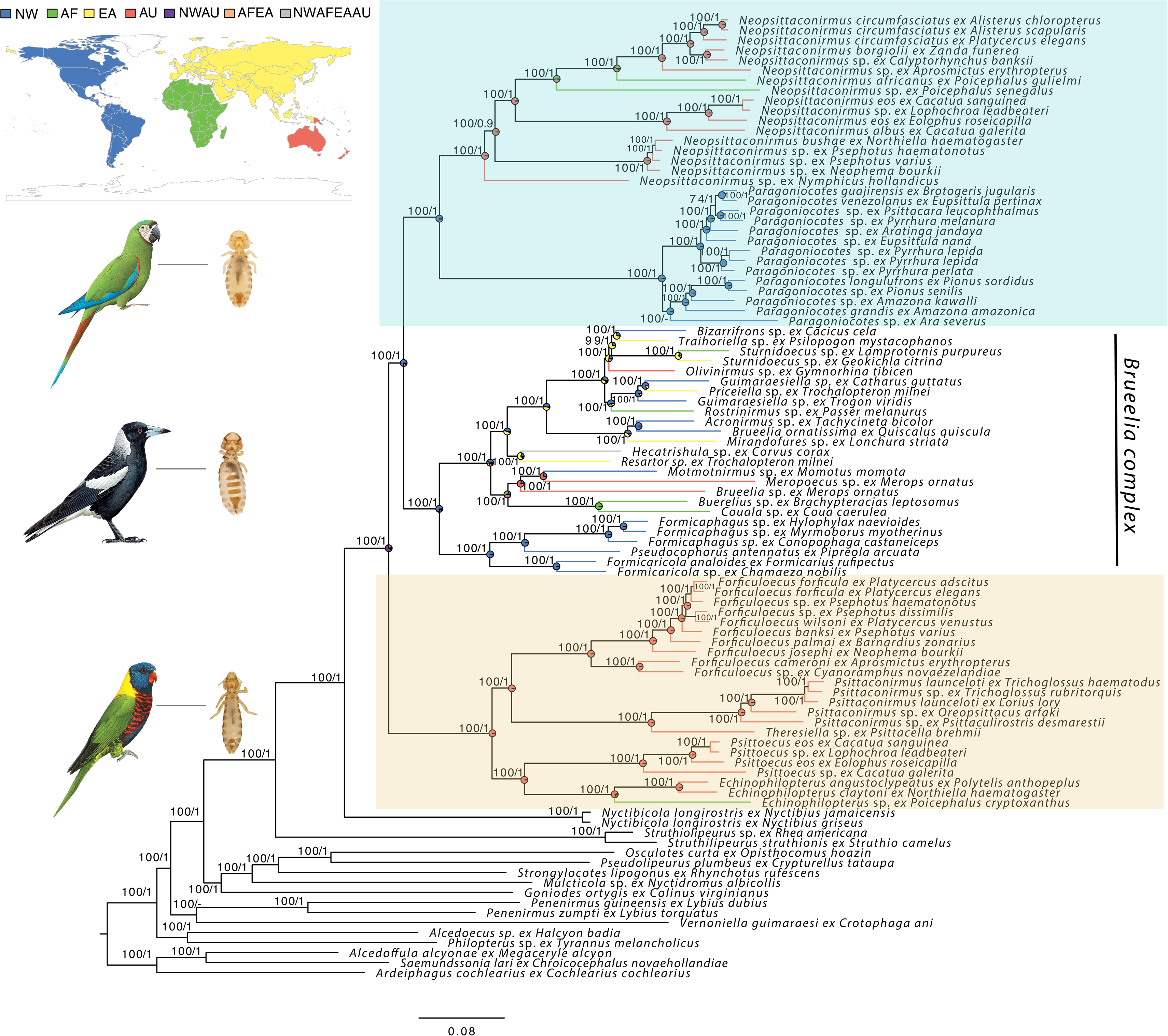
Phylogeny from partitioned concatenated analysis of 2,395 target nuclear gene orthologs for parrot feather lice and relatives. Numbers on branches indicate support from ultrafast bootstrap replicates/coalescent local posterior probability. The two major clades of parrot lice and the *Brueelia* complex are highlighted with images of representative taxa and their hosts. Pie charts at nodes indicate relative likelihoods for ancestral distributions inferred by BioGeoBears under the DEC+J model. Colors correspond to major biogeographical regions indicated by map inset. Bird illustrations reprinted by permission ©Lynx Edicions.

Both trees indicated that parrot feather lice as a group are paraphyletic, falling into two distinct clades (Figure 1). One of these clades contained all samples of *Neopsittaconirmus* and *Paragoniocotes*, and this clade was sister to the *Brueelia*-complex. The second clade contained the parrot louse genera *Forficuloecus*, *Psittaconirmus*, *Theresiella*, *Psittoecus*, and *Echinophilopterus*. This second clade was sister to the first clade plus the *Brueelia*-complex. Within this second clade, *Echinophilopterus* and *Psittoecus* were sister taxa, and together these two genera were sister to a clade comprising *Theresiella*, *Psittaconirmus*, and *Forficuloecus*. Among parrot lice, all genera were reconstructed as monophyletic, albeit the monophyly of *Theresiella* could not be tested because it was sampled by only a single species. As in prior studies, the genus *Nyctibicola* from potoos (Nyctibiidae) was sister to the parrot louse and *Brueelia*-complex clade.

This phylogenetic arrangement of genera indicates that parrot louse ecomorphs evolved multiple times. For example, the two wing louse genera, *Psittaconirmus* and *Neopsittaconirmus*, are distantly related being in different clades. Similarly, the two body louse genera, *Psittoecus* and *Paragoniocotes*, are also in distinct clades. The three head louse genera, *Echinophilopterus*, *Theresiella*, and *Forficuloecus*, are also not sister taxa, even though they are all in the same clade. However, beyond these major ecomorph transitions between genera, the fact that the genera are monophyletic indicates that there are not further ecomorphological changes among parrot lice.

### Mitochondrial COI divergences and species boundaries

Within each of the two parrot louse clades, most pairwise uncorrected mitochondrial divergences for the COI gene far exceeded 10% (Figure 2). In some cases, this divergence was between individuals of what would otherwise be considered the same species, indicating potential cryptic species. For example, within *Neopsittaconirmus circumfasciatus*, the louse from *Platycercus elegans* was 15-16% divergent from those on the two *Alisterus* host species. Within *Neopsittaconirmus eos*, the louse from *Eolophus rosiecapilla* was 16.0% divergent from those on *Cacatua* hosts. Finally, within *Psittoecus eos*, the louse from *Cacatua sanguinea*, was 19.6% divergent from those individuals on *Lophochroa leadbeateri* and *Eolophus roseicapillus*.

**Figure 2.**
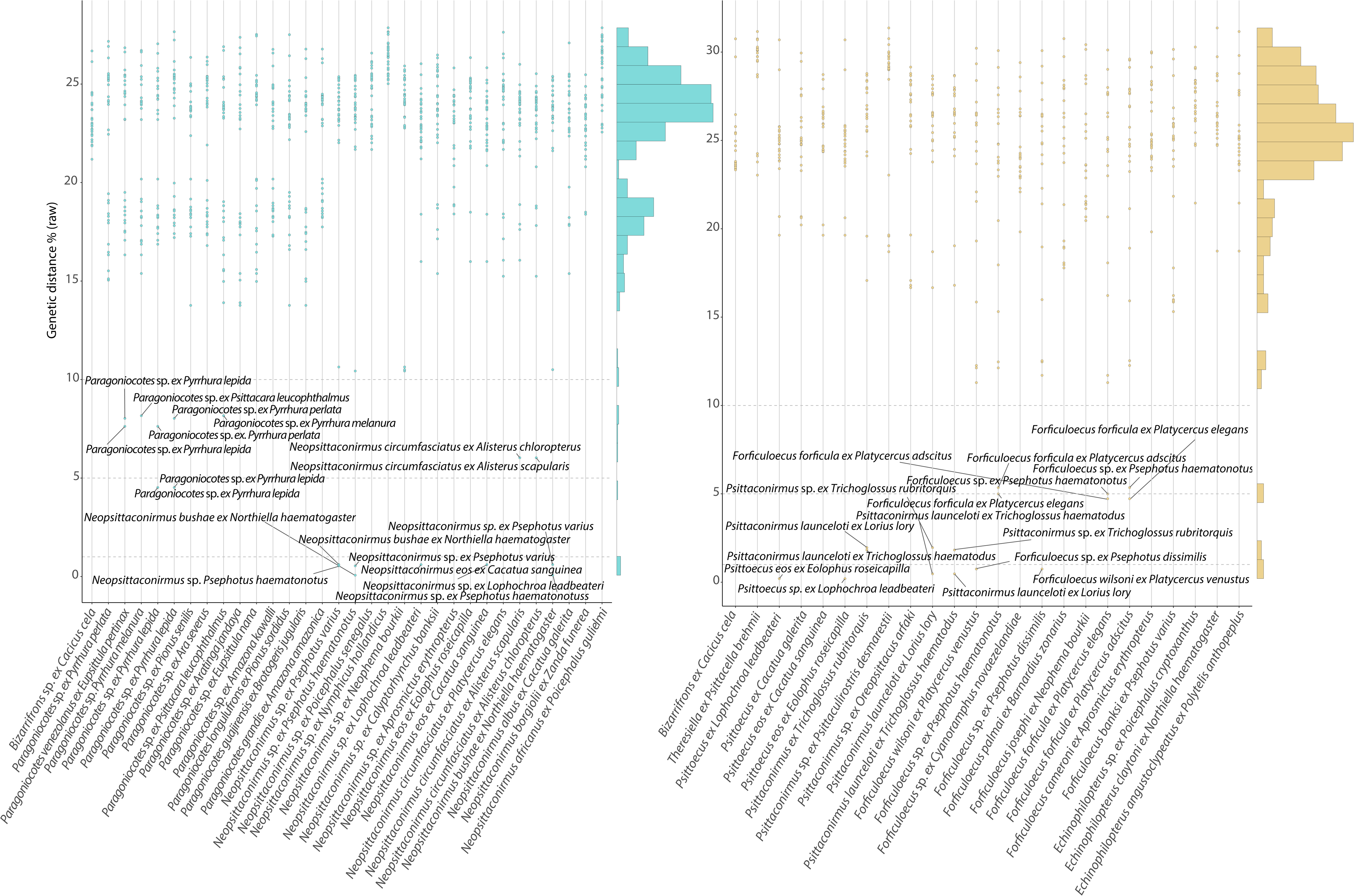
Pairwise uncorrected mitochondrial COI divergences (y-axis) for the two parrot louse clades. Histograms indicate relative frequency of divergences among all pairwise comparisons.

There were also a few comparisons in each clade that were less than around 2%, indicating likely conspecific samples. In particular, samples of *Neopsittaconirmus* from three different Australian parrots (*Northiella haematogaster*, *Psephotus varius*, and *P. haematonotus*) all showed less than 1% COI sequence divergence from each other, indicating they are all likely the same species, *Neopsittaconirmus bushae*. Similarly, individuals of *Neopsittaconirmus* from two different Australian cockatoos (*Cacatua sanguinea* and *Lophochroa leadbeateri*) were only 0.61% divergent from each other. Two individuals of *Psittoecus* had a different pattern of host sharing, with individuals from *Lophochroa leadbeateri* and *Eolophus roseicapillus* being nearly identical (0.20% divergence) to each other. The three samples of *Psittaconirmus* from lorikeets (*Trichoglossus hamatodus*, *T. rubritorquis*, and *Lorius lory*), also showed very little divergence (all less than 2%) from each other. Finally, samples of *Forficuloecus* from two Australian parrots (*Platycerus venustus* and *Psephotus dissimilis*) were also nearly identical (0.75%), again likely indicating a louse species (*Forficuloecus wilsoni*) that is not completely host specific. Interestingly, lice from other hosts in the *Platycercus* species complex (Joseph et al. 2008), are much more divergent from *F. wilsoni* on *Platycercus venustus* (> 11.0% COI).

In a few cases, pairwise comparisons produced COI divergences near our *a priori* 5% cutoff for distinguishing species on the basis of genetic divergence. The comparison within *Paragoniocotes* (4.52%) involved lice from two subspecies of the Pearly Parakeet (*Pyrrhura lepida*; ssp. *anerythra* and *lepida*), and this likely represents respective population divergence in the lice. The case in *Neopsittaconirmus* (6.03%) involved lice from two congeneric species of king parrots (*Alisterus scapularis* and *chloropterus*) from Australia and New Guinea, respectively, and likely represents codivergence with their hosts, supporting the idea that 5% may be an appropriate cutoff for diagnosing species of *Paragoniocotes* and *Neopsittaconirmus*. In the case of *Forficuloecus*, three individuals of *Forficuloecus forficula* (from hosts *Psephotus haematonotus*, *Platycercus adscitus*, and *Platycercus elegans*) had COI divergences near 5%. However, the divergence between the lice from the two *Platycercus* was 4.72%, while the divergences between the louse from *Psephotus haematonotus* and those on *Platycercus* were 5% or more, suggesting these lice may be on the border of being cryptic species.

### Species level phylogeny

Given our taxon sampling, we were also able to recover broad patterns of species level diversification within some of the more densely sampled genera. Generally, results from COI divergences corresponded to the structure of the tree from nuclear orthologs (Figure 1). Within *Paragoniocotes*, which is restricted to the New World, lice from smaller bodied parakeets (*Pyrrhura*, *Aratinga*, *Eupsittula*, *Brotogeris*), clustered together to the exclusion of larger bodied parrots (*Amazona*, *Pionus*) and macaws (*Ara*), the latter two forming either a clade (concatenated tree) or grade (coalescent tree). Within these groups of lice on small and large bodied hosts, there were not additional clades of lice exclusive to a single host genus.

Within *Neopsittaconirmus*, there was not clear partitioning by host group. Lice from cockatoos occurred on three separate branches. Interestingly lice from white cockatoos and black cockatoos were separated, perhaps related to their cryptic coloration in adaptation to host feather color (Bush et al. 2010). The lice from African parrots (two species of *Poicephalus*) were also embedded within an otherwise Australasian clade. Like *Paragoniocotes*, there did seem to be some size partitioning with lice from small-bodied hosts at the base of the tree, and those from larger bodied hosts forming a separate clade.

The genus *Forficuloecus* was divided into two highly divergent clades. One with two species (*Forficuloecus pilgrimi* and *cameroni*) occurring on the Red-fronted Parrot (*Cyanoramphus novaezelandiae*) from New Zealand and the Red-winged Parrot (*Aprosmictus erythropterus*) from Australia, respectively. This clade was sister to the other major clade which contained lice from a variety of small and mid-sized parrots from Australia.

The genus *Psittaconirmus* had a clade of lice from comparatively large-bodied lorikeets (*Trichoglossus* and *Lorius*), embedded within lice from small-bodied lorikeets and fig parrots (*Oreopsittacus* and *Psittaculirostris*). This clade is entirely Australasian and sister to another Australiasian genus, *Theresiella*, represented in our study by a specimen from a tiger-parrot (*Psittacella*) from New Guinea.

The genera *Psittoecus* and *Echinophilopterus* were found to be sister taxa, and each has only a small number of described species (five and seven respectively). The genus *Psittoecus* is exclusive to cockatoos, but there was no obvious partitioning by host size. The two species of *Echinophilopterus* from Australia were sister taxa, with the sample from an African parrot being more divergent.

### Biogeography

Within parrot lice there was generally strong biogeographic signal. The model favored by AIC in BioGeoBears was DEC+J (dispersal-extinction-colonization, plus jump dispersal). Under this model (Figure 1), the ancestral distribution of the *Neopsittaconirmus* + *Paragoniocotes* clade was ambiguous, with nearly equal probabilities of being either New World or Australasian in distribution. The ancestral distribution of *Paragoniocotes* was inferred to be New World, which is not surprising given its entirely New World distribution. The ancestral distribution of *Neopsittconirmus* was inferred to be Australasian with up to two dispersal events to Africa. Similarly, the ancestral distribution of the *Forficuloecus* + *Psittaconirmus* + *Theresiella* + *Psittoecus* + *Echinophilopterus* clade was inferred to be Australasian with a dispersal event into Africa.

## Discussion

Phylogenomic analyses of 2,395 target nuclear ortholog genes for the feather lice of parrots and related taxa produced a well resolved and strongly supported tree. Parrot lice fell into two major clades that formed a paraphyletic grade with respect to the *Brueelia*-complex. One of these clades was restricted to the Old World and contained two genera of head lice (*Echinophilopterus* and *Forficuloecus*), body lice from cockatoos (*Psittoecus*), and two genera of wing lice (*Psittaconirmus* and *Theresiella*) from lorikeets and other Australiasian parrots. The second major clade comprised a genus of Old World wing lice (*Neopsittaconirmus*) sister to New World body lice (*Paragoniocotes*). In all cases, we found strong support of monophyly of genera that were sampled by more than one species, indicating that current generic limits correspond to evolutionary relationships.

Previous data sets that have sampled both of these major clades have been unclear on whether these two major clades of parrot lice form a monophyletic group (de Moya et al. 2019, de Moya 2022) or not (Johnson et al. 2012). These prior studies only sample one or a few species per major clade, and here we dramatically increase the taxon sampling to include over 20 species per clade plus extensive outgroup sampling, particularly in the *Brueelia*-complex. With this increased sampling, we find 100% bootstrap (concatenated) and 1.0 local posterior probability (coalescent) for paraphyly of parrot lice with respect to the *Brueelia*-complex. Parrots (Psittaciformes) are sister to songbirds (Passeriformes) (Jarvis et al. 2014, Prum et al. 2015), and given that the predominant hosts of lice with in the *Brueelia*-complex occur on songbirds, this may be an indication of ancient codivergence. However, given the paraphyly of parrot lice, the situation may be more complex.

Among the major lineages of parrot lice, both biogeography and host association appeared to provide the overall structure to the phylogenetic tree. Major clades and genera are restricted to particular biogeographic regions. In general, there were few transitions between biogeographic regions across the louse tree. However, parrot phylogeny itself is also related to biogeography, with all New World parrots forming a clade (Wright et al. 2008). Thus, biogeography and host association are intertwined and cannot be completely disentangled as explanations for the structure of the louse tree.

Within genera of parrot lice, some of the phylogenetic structure appeared to be related to host body size. Species on smaller and larger bodied hosts tended to cluster together respectively. Again, body size is also partly related to host phylogeny (Wright et al. 2008), so these two factors could be difficult to disentangle, but it does suggest that any host switching that does occur might be limited to hosts of similar body size.

Overall, our findings offer a robust foundational framework for understanding the phylogeny of parrot lice. They indicate that current generic limits are well supported. However, we found evidence for cryptic genetic variation within species of lice, particularly those that occur on multiple host species. Future taxonomic work should focus on more clearly defining species boundaries, integrating morphological revision with molecular analyses.

## Acknowledgements

We thank N. Aristizábal, B. Benz, S. E. Bush, T. Chesser, D. H. Clayton, L. Cueto, R. Empson, R. Faucett, T. D. Galloway, A. Gouvea, E. H. Kuschel, D. Lane, I. Mason, R. Moyle, B. O’Shea, R. Palma, R. Palmer, V. Q. Piacentini, A. Porzecanski, V. Smith, S. Sontshugen, J. D. Weckstein, R. Wilson, and J. Wombey for assistance in obtaining samples for this study. A. Hernandez and C. Wright at the University of Illinois Roy J. Carver Biotechnology Center provided assistance with Illumina genome sequencing. We thank K. K. O. Walden for assistance with submission of raw reads to NCBI SRA. Funding was provided by U.S. NSF DEB-1239788, DEB-1925487, and DEB-1926919 to K.P.J. and European Commission grant H2020-MSCA-IF-2019 (INTROSYM: 8865532) to J.D.

## Data availability

Data generated in this study are available at Figshare (https://figshare.com/s/4f33cbe035e9da9b846c; 10.6084/m9.figshare.23807646).

## Notes

https://doi.org/10.6084/m9.figshare.23807646

### Competing Interest Statement

The authors have declared no competing interest.

