## Supplementary material for "Phylogenomics and biogeography of the feather lice (Phthiraptera: Ischnocera) of parrots": Table S1

**Supplemental Table S1. Calibration points used in molecular dating analysis**

| **Calibration Node*** | **Date** |
| --- | --- |
| *Psittaconirmus sp.* | 13.5 – 18.5 mya |
| *Paragoniocotes longulufrons* | 3.0– 6.5 mya |
| *Neopsittaconirmus borgiolii* | 10.0 – 15.0 mya |
| *Neopsittaconirmus circumfasciatus* | 2.0 mya |
| *Paragoniocotes sp.* | 0.5 mya |

*Indicates date range for node uniting taxon indicated with its sister taxon (from Figure 1)
